# The p53 transactivation domain 1-dependent response to acute DNA damage in endothelial cells protects against radiation-induced cardiac injury

**DOI:** 10.1101/2021.12.04.471191

**Authors:** Hsuan-Cheng Kuo, Lixia Luo, Yan Ma, Nerissa T. Williams, Lorraine da Silva Campos, Laura D. Attardi, Chang-Lung Lee, David G. Kirsch

## Abstract

Thoracic radiation therapy can cause endothelial injury in the heart, leading to cardiac dysfunction and heart failure. Although it has been demonstrated that the tumor suppressor p53 functions in endothelial cells to prevent the development of radiation-induced myocardial injury, the key mechanism(s) by which p53 regulates the radiosensitivity of cardiac endothelial cells is not completely understood. Here, we utilized genetically engineered mice that express mutations in p53 transactivation domain 1 (TAD1) (p53^25,26^) or mutations in p53 TAD1 and TAD2 (p53^25,26,53,54^) specifically in endothelial cells to study the p53 transcriptional program that protects cardiac endothelial cells from ionizing radiation *in vivo*. p53^25,26,53,54^ loses the ability to drive transactivation of p53 target genes after irradiation while p53^25,26^ can induce transcription of a group of non-canonical p53 target genes, but not the majority of classic radiation-induced p53 targets critical for p53-mediated cell cycle arrest and apoptosis. After 12 Gy whole-heart irradiation, we found that both p53^25,26^ and p53^25,26,53,54^ sensitized mice to radiation-induced cardiac injury, in contrast to wild-type p53. Histopathological examination suggested that mutation of TAD1 contributes to myocardial necrosis after whole-heart irradiation, while mutation of both TAD1 and TAD2 abolishes the ability of p53 to prevent radiation-induced heart disease. Taken together, our results show that the transcriptional program downstream of p53 TAD1, which activates the acute DNA damage response after irradiation, is necessary to protect cardiac endothelial cells from radiation injury *in vivo*.

## Introduction

Understanding the mechanisms regulating radiation-induced normal tissue injury can inform the design of improved approaches to treating cancer with radiation therapy. A better understanding of the mechanisms of normal tissue injury from radiation could also facilitate the development of therapeutic interventions for potential radiation accidents or disasters. The heart can be exposed to radiation therapy during treatment for breast cancers, lymphomas, lung cancers, or other tumors. Although the heart has been considered a relatively radioresistant organ, the cardiovascular system displays various pathologies following radiation exposure (Spetz, Moslehi, & Sarosiek, 2018) and recent studies suggest that a mean radiation dose of 5 Gy to the heart can cause subsequent heart disease (Darby et al., 2013). A recently published analysis of 972 women who received radiation therapy (RT) for breast cancer and had no pre-existing cardiovascular disease showed that women who received left-sided RT had a higher risk of coronary artery disease (CAD) than those who received right-sided RT (HR: 2.5) (Bates, Chen, & Constine, 2021; Carlson et al., 2021). The 27.5-year cumulative incidence of CAD was 10.5% in women treated with left-sided RT, which was significantly higher than the incidence in right-sided RT-treated women (5.8%) (Carlson et al., 2021). Moreover, cancer patients who receive thoracic radiotherapy have a 1.5 to 3 times higher risk of a fatal cardiac event than those who do not receive radiotherapy (Spetz et al., 2018). Some of the atomic bomb survivors also developed an increased risk of cardiovascular disease years after the radiation exposure (Boerma et al., 2016). Cardiovascular injury after radiation injury may be manifested as myocardial necrosis, vascular and valvular damage, and pericardial inflammation and fibrosis (Spetz et al., 2018). Radiation exposure to the heart can lead to systolic dysfunction, coronary artery disease, and heart failure (Boerma et al., 2016; Lee et al., 2012; Spetz et al., 2018), which may be a consequence of endothelial cell disruption and dysfunction (Korpela & Liu, 2014). Endothelial cells are a critical cell type in the heart that responds to radiation by undergoing cell death, senescence, or alterations of cytokine production (Korpela & Liu, 2014). These responses of endothelial cells to radiation lead to the disruption of microvasculature, hypoxia, myocardial necrosis, and tissue remodeling including myocardial fibrosis that can compromise cardiac contraction and cause arrhythmias and even death (Wang et al., 2020).

The DNA damage response (DDR) leads to many sequelae after exposure to ionizing radiation (Jackson & Bartek, 2009). One key protein that responds to DNA damage is the tumor suppressor p53 (Kastenhuber & Lowe, 2017). Acute DNA damage activates p53 to induce apoptosis and/or cell cycle arrest in a cell type-dependent manner (Bieging, Mello, & Attardi, 2014; Kastenhuber & Lowe, 2017). We have shown that p53 functions in cardiac endothelial cells to prevent the development of radiation-induced myocardial injury (Lee et al., 2012). Deletion of both alleles of *p53* in endothelial cells by Cre recombinase expressed from either the *Tie2* or vascular endothelial cadherin (*VE*) promoter exacerbated cardiac dysfunction after 12 Gy whole-heart irradiation (WHI) (Lee et al., 2012). However, given that p53 controls a variety of signaling pathways including apoptosis, cell cycle arrest, DNA repair and metabolism, the key mechanism(s) by which p53 protects cardiac endothelial cells from radiation injury *in vivo* remains incompletely understood.

Following ionizing radiation, p53 functions as a transcription factor to activate the expression of hundreds of genes (Boutelle & Attardi, 2021; Kaiser & Attardi, 2018). p53-mediated target gene transcription requires two distinct domains in the N-terminus, namely transactivation domain 1 (TAD1) and 2 (TAD2) (Bieging & Attardi, 2012; Bieging et al., 2014). TAD1 and TAD2 work coordinately to interact with several proteins that facilitate the initiation of RNA polymerase II-mediated gene transcription or other transcription-related events (Bieging & Attardi, 2012). Specific amino acid residues within p53 TADs are critical for transactivation function. Mutation of the 25^th^ and 26^th^ residues of mouse p53 (22^nd^ and 23^rd^ residues in human p53) compromises TAD1 function on many p53 target genes while mutation of the 53^rd^ and 54^th^ residues (53^rd^ and 54^th^ residues in human p53) disrupts TAD2 function in the context of TAD1 mutation, leading to a transactivation defective mutant (Bieging & Attardi, 2012). Mutating p53 at these amino acids to hydrophobic residues alters transactivation, which provides a genetic tool to investigate the role of p53-induced gene transcription in tumor suppression and other functions *in vivo*. In *p53^25,26/-^* cells, the mutated TAD1 domain is unable to drive the DNA damage-induced transcription of canonical p53 target genes such as *PUMA* and the cyclin-dependent kinase inhibitor *p21*, although the TAD2 domain in these cells remains functional to induce transcription of a small portion of non-canonical p53 target genes (Brady et al., 2011). In contrast, *p53^25,26,53,54/-^* cells that harbor mutations in both TAD1 and TAD2 domains are deficient in activating all transcriptional targets of p53 and therefore resemble p53 null cells (Brady et al., 2011). These mutant alleles provide a sophisticated genetic tool to investigate the mechanism(s) by which p53 functions in endothelial cells to regulate radiation-induced heart disease by separating transactivation domain-dependent events into ones that require functional TAD1 and TAD2 with those that require TAD2 alone. Here, we performed experiments using mice with endothelial cells that lack p53 expression or express different TAD mutants to determine the role of p53 transcriptional targets induced by TAD1 (acute DNA damage response) vs. TAD1 and TAD2 (all p53 targets) in controlling the radiation response of cardiac endothelial cells *in vivo*.

## Materials and methods

### Mice

All procedures with mice were approved by the Institutional Animal Care and Use Committee (IACUC) of Duke University. All mouse strains used in this study have been described before, including *VE-Cadherin-Cre* (*VE-Cre*), *p53^flox^, p53^LSL-25,26^*, and *p53^LSL-25,26,53,54^* mice (Alva et al., 2006; Brady et al., 2011; Marino, Vooijs, van Der Gulden, Jonkers, & Berns, 2000). The *VE-Cre* mice were originally obtained from the Jackson Laboratory and then bred at Duke University. The *p53^flox^* mice were originally provided by A. Berns (Netherlands Cancer Institute, Amsterdam, the Netherlands) and then bred at Duke University. Mice were at least 8 weeks old at the time of irradiation. Both sexes of mice were used. Comparisons were made using littermate controls to minimize genetic differences as these experiments were performed on mixed genetic backgrounds.

### Radiation treatment

For whole heart irradiation (WHI) treatments (Lee et al., 2012), a small-field image-guided irradiator, X-RAD 225Cx (Precision X-Ray Inc.), was used with fluoroscopy. Radiation treatments were performed with parallel-opposed anterior and posterior fields as we previously described. A cone was inserted as a collimator to create a 15mm circular radiation field at the isocenter where the heart was localized.

### EF5 hypoxia detection

10mM EF5 (Millipore Sigma) solution was prepared according to the product data sheet protocol. The solution was administered through intraperitoneal injection at 26.5μL/g body weight to mice. Hearts of the injected mice were harvested 3 hours after the injection, and then fixed with 4% paraformaldehyde for frozen section preparations. The antibody Anti-EF5, clone ELK3-51 Cyanine 3 conjugate (Millipore Sigma) was used to detect EF5, and Isolectin GS-IB_4_ Alexa Fluor 647 conjugate (Invitrogen) was used to detect vasculature.

### Histological Analyses

Tissues were fixed with 10% neutralized formalin for paraffin-embedded tissue specimens. Slides were stained with H&E or Masson’s trichrome staining. For immunohistochemistry (IHC), de-paraffinized slides underwent 3% hydrogen peroxide treatment, antigen retrieval with citrate-based solution, and blocking by serum. Slides were incubated with primary antibody against p53 protein (CM5; Leica Biosystems) and then with secondary antibody. The VECTASTAIN Elite ABC system (Vector Laboratories) was applied, and 3,3’-diaminobenzidine (DAB) was used as chromogen.

For frozen tissue specimens, harvested tissues were fixed with 4% paraformaldehyde in phosphate-buffered saline (PBS) at 4°C, and then placed in 30% sucrose in PBS overnight at 4°C. Processed tissues were then frozen in cryomolds of OCT compound (Sakura Finetek) and stored at −80°C. The heart of each mouse was cut in half before placing into a mold with the cut section facing the sectioning plane.

### Statistics

The Kaplan-Meier method was used to analyze the development of radiation-induced heart disease. Statistical comparison was performed using log-rank test. GraphPad Prism (GraphPad Software) was used for the statistics.

## Results

To investigate the role of different p53 TADs in regulating the radiosensitivity of cardiac endothelial cells *in vivo*, VE-Cre was used to recombine the floxed allele of *p53* (*p53^FL^*) and remove the STOP cassette in the *p53* TAD mutant alleles (**Figure 1A**). We previously reported the distribution of VE-Cre activity within the epimyocardium and midmyocardium of the heart as measured by Cre-mediated recombination (Lee et al., 2012). The functional status of p53 in different transgenic mice used in this study is summarized in **Figure 1B**.

**FIG. 1.**
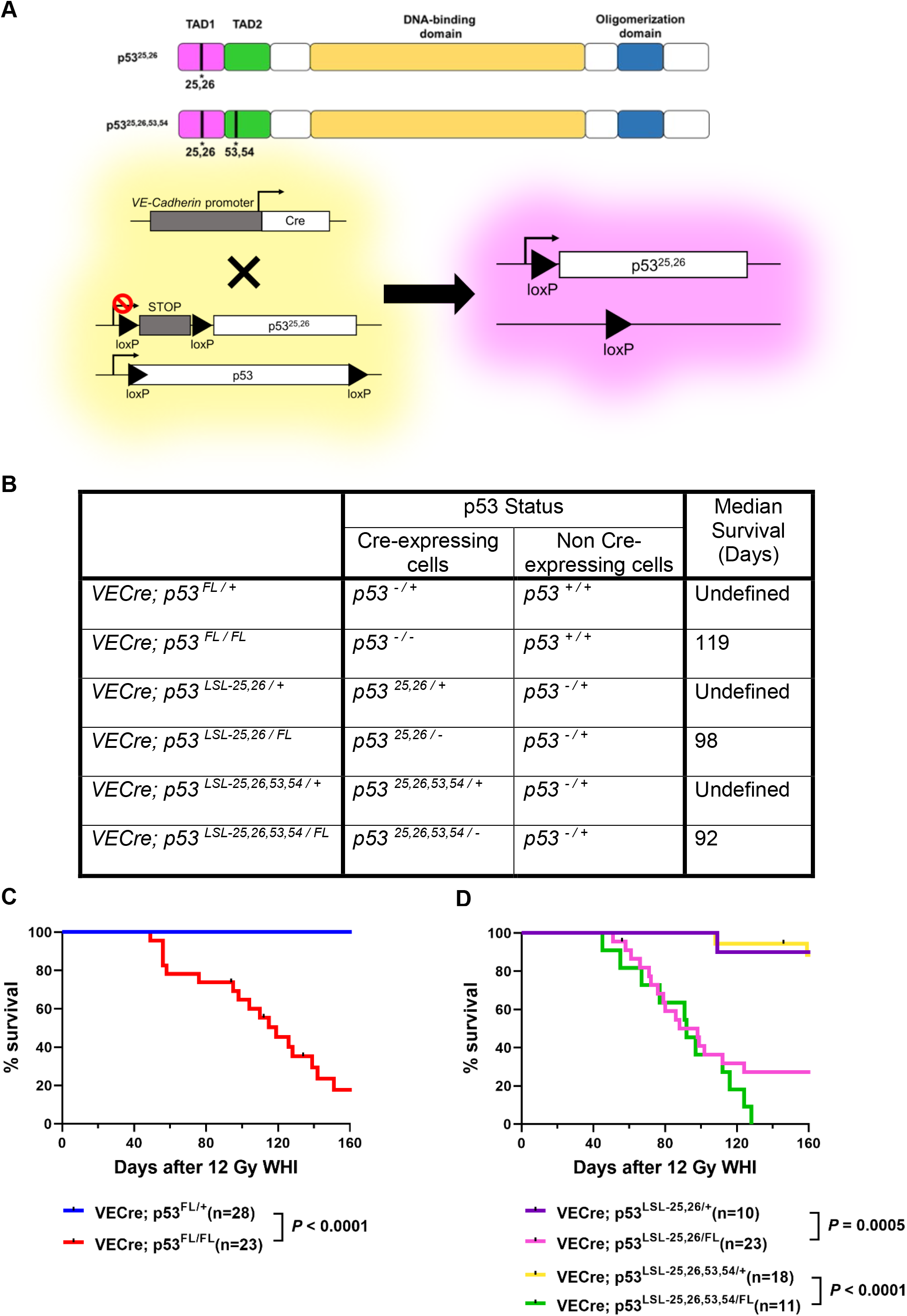
p53 TAD1 in endothelial cells protects against radiation-induced cardiac injury *in vivo*. Panel A: A schematic showing the domain composition of p53 protein and the mutation sites of p53^25,26^ and p53^25,26,53,54^ (Brady et al., 2011; Jiang et al., 2015) (top) and a schematic showing VE-Cre-mediated recombination of the STOP cassette to express a p53 TAD mutant in the endothelial cell (bottom). Panel B: A table of the mouse genotypes used in this study. The p53 status of Cre-expressing and non Cre-expressing cells in each genotype is shown. Mice were treated with 12 Gy whole-heart irradiation (WHI). The rightmost column shows the median survival of each genotype. Panel C & D: Mice received 12 Gy WHI and were followed for the development of radiation-induced cardiac injury. The Kaplan-Meier curves of each genotype are shown in different colors. P-values (*P*) were calculated using log-rank test.

All mice that expressed various *p53* mutant alleles specifically in the endothelial cells were treated with 12 Gy WHI. Previously, we found that *VE-Cre; p53^FL/-^* mice developed heart failure within 40 to 130 days after WHI (Lee et al., 2012). Here, we found similar results in which *VE-Cre; p53^FL/FL^* mice that lack p53 in the endothelial cells were significantly more susceptible to the development of radiation-induced cardiac injury compared with *VE-Cre; p53^FL/+^* mice that retain one allele of *p53* in the endothelial cells (**Figure 1C**). These findings demonstrate that 12 Gy WHI reproducibly induces heart disease in mice lacking p53 in endothelial cells. Similar to *VE-Cre; p53^FL/FL^* mice, *VE-Cre; p53^LSL-25,26,53,54/FL^* mice succumbed to radiation-induced cardiac injury within 40 to 130 days after 12 Gy WHI (**Figure 1D**), while *VE-Cre; p53^LSL-25,26,53,54/+^* mice that retain one wild type allele of *p53* in the endothelial cells were not sensitized to radiation-induced cardiac injury (**Figure 1D**). In addition, *VE-Cre; p53^LSL-25,26/FL^* mice were also significantly more susceptible to the development of radiation-induced cardiac injury after 12 Gy WHI compared with *VE-Cre; p53^LSL-25,26/+^* mice that retain one wild type allele of *p53* in the endothelial cells. Notably, the latency of radiation-induced cardiac injury was similar between *VE-Cre; p53^LSL-25,26/FL^* mice and *VE-Cre; p53^LSL-25,26,53,54/FL^* mice (**Figure 1D**). Additionally, we found strong p53 protein staining in cardiac endothelial cells of *VE-Cre; p53^LSL-25,26/+^* mice and *VE-Cre; p53^LSL-25,26,53,54/+^* mice because TAD mutations lead to the accumulation of p53 protein in cells (**Supplementary figure 1**) at least in part because these mutants compromise the p53-MDM2 interaction (Van Nostrand et al., 2014). Nevertheless, the accumulation of p53 TAD mutant proteins did not impact the ability of the wild type *p53* allele to prevent the development of radiation-induced cardiac injury, and it might have potentially stimulated wild type p53 activity (Van Nostrand et al., 2014). Collectively, our results from mice that have defects in p53 TAD1 (*VE-Cre; p53^LSL-25,26/FL^*) or p53 TAD1/TAD2 (*VE-Cre; p53^LSL-25,26,53,54/FL^*) in endothelial cells indicate that transcriptional targets of p53, especially the ones induced by p53 TAD1, are necessary to protect cardiac endothelial cells against radiation injury.

To investigate the causes leading to the development of cardiac injury in the p53 TAD mutants, we collected the hearts from different genotypes at day 55 following 12 Gy WHI. We found that both *VE-Cre; p53^FL/FL^* and *VE-Cre; p53^LSL-25,26,53,54/FL^* mice displayed extensive myocardial necrosis, while necrosis in the myocardium of *VE-Cre; p53^LSL-25,26/FL^* mice was instead focal to multifocal (**Figure 2**). In contrast, *VE-Cre; p53^FL/+^, VE-Cre; p53^LSL-25,26/+^*, and *VE-Cre; p53^LSL-25,26,53,53/+^* mice that retained a *p53* wild type allele in endothelial cells displayed no or minimal myocardial necrosis (**Figure 2**).

**FIG. 2.**
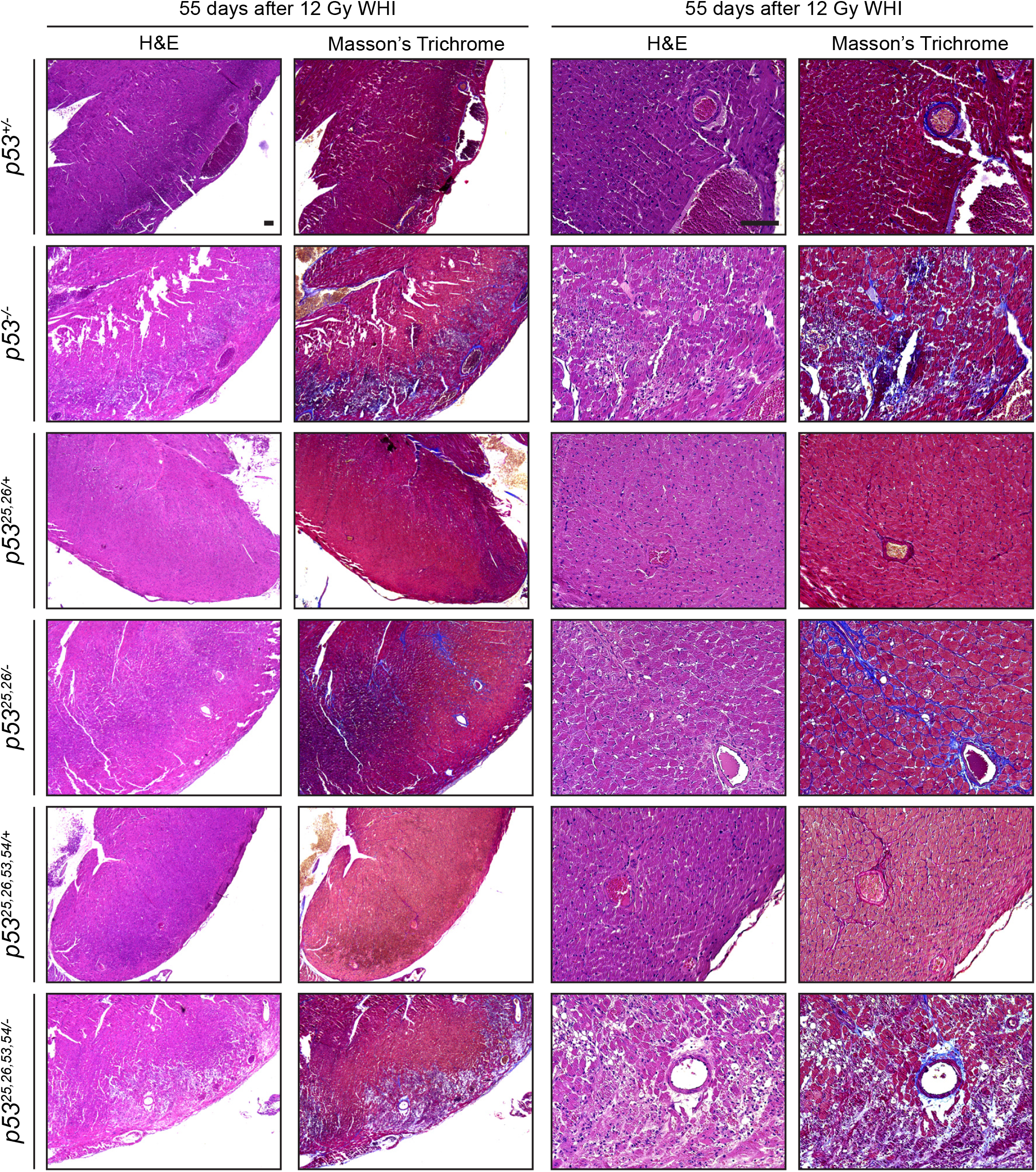
p53 TAD functions in endothelial cells regulate myocardial necrosis and fibrosis after WHI. Hearts were harvested from mice of different genotypes at day 55 after 12 Gy WHI. Hematoxylin and eosin (H&E) staining and Masson’s trichrome staining were performed. Representative images of different genotypes are shown at two magnifications. The p53 status of Cre-expressing cells from each genotype is labeled at the left. Scale bars, 100μm.

We also examined fibrosis in the irradiated hearts by Masson’s trichrome staining. Similar to the H&E results, both *VE-Cre; p53^FL/FL^* and *VE-Cre; p53^LSL-25,26,53,54/FL^* showed prominent pathological changes in the myocardium, which included tissue fibrosis and disruption of the myocardium (**Figure 2**). The disrupted regions of the myocardium were replaced with fibrosis and they corresponded to the necrotic regions shown by H&E. Different levels of interstitial fibrosis were observed across the tissue sections. Similar to what we found with H&E staining, *VE-Cre; p53^LSL-25,26/FL^* samples showed intermediate results with some of the samples showing minimal fibrosis and others showing more profound fibrosis (**Figure 2**). Taken together, loss of p53 transactivation (p53^25,26,53,54^) led to multifocal myocardial necrosis, suggesting that the function of the TADs of p53 in the endothelial cells are essential to prevent radiation-induced myocardial necrosis. Aberrant p53 transactivation capability (p53^25,26^) did not cause the same amount of profound myocardial necrosis, but still led to the development of cardiac injury that debilitated the animal, highlighting the critical importance of p53-mediated transactivation of TAD1 target genes. There were almost no disrupted myocardial regions in the *VE-Cre; p53^FL/+^* samples 55 days after 12 Gy WHI. Similarly, there was no to minimal fibrosis in the *VE-Cre; p53^FL/+^, VE-Cre; p53^LSL-25,26/+^*, and the *VE-Cre; p53^LSL-25,26,53,53/+^* samples (**Figure 2**). These results suggest that the presence of wild type p53 in endothelial cells prevented the development of radiation-induced tissue fibrosis.

To understand if dysregulated endothelial response led to myocardial necrosis through tissue hypoxia, we utilized EF5 to label hypoxic regions as we previously reported (Lee et al., 2012). As expected, *VE-Cre; p53^FL/FL^* samples showed multiple regions of EF5-labeled fluorescence signal while *VE-Cre; p53^FL/+^* samples showed none or minimal lesions (**Figure 3**). The *VE-Cre; p53^LSL-25,26,53,54/FL^* and the *VE-Cre; p53^LSL-25,26/FL^* samples showed EF5-labeled hypoxic regions and/or disorganized vasculature detected by the glycoprotein GS-IB_4_, though the *VE-Cre; p53^LSL-25,26/FL^* samples were not as prominent. (**Figure 3**). This suggests that in endothelial cells the non-canonical p53 targets from TAD2 may limit the extent of radiation-induced myocardial necrosis, possibly by preventing hypoxia. In contrast, there were no or much fewer hypoxic regions and/or disorganized vasculature in the *VE-Cre; p53^FL/+^, VE-Cre; p53^LSL-25,26/+^*, and the *VE-Cre; p53^LSL-25,26,53,53/+^* samples from mice that retain a wild type *p53* allele in endothelial cells (**Figure 3**). The results from the histopathology indicate that extensive radiation-induced myocardial necrosis correlates with prominent hypoxia and/or disorganized vasculature. Taken together, this study provides evidence that inactivation of p53 TAD1 in endothelial cells is sufficient to sensitize mice to radiation-induced cardiac injury, while more severe damage to the myocardium occurs in mice where both p53 TAD1 and TAD2 are inactivated.

**FIG. 3.**
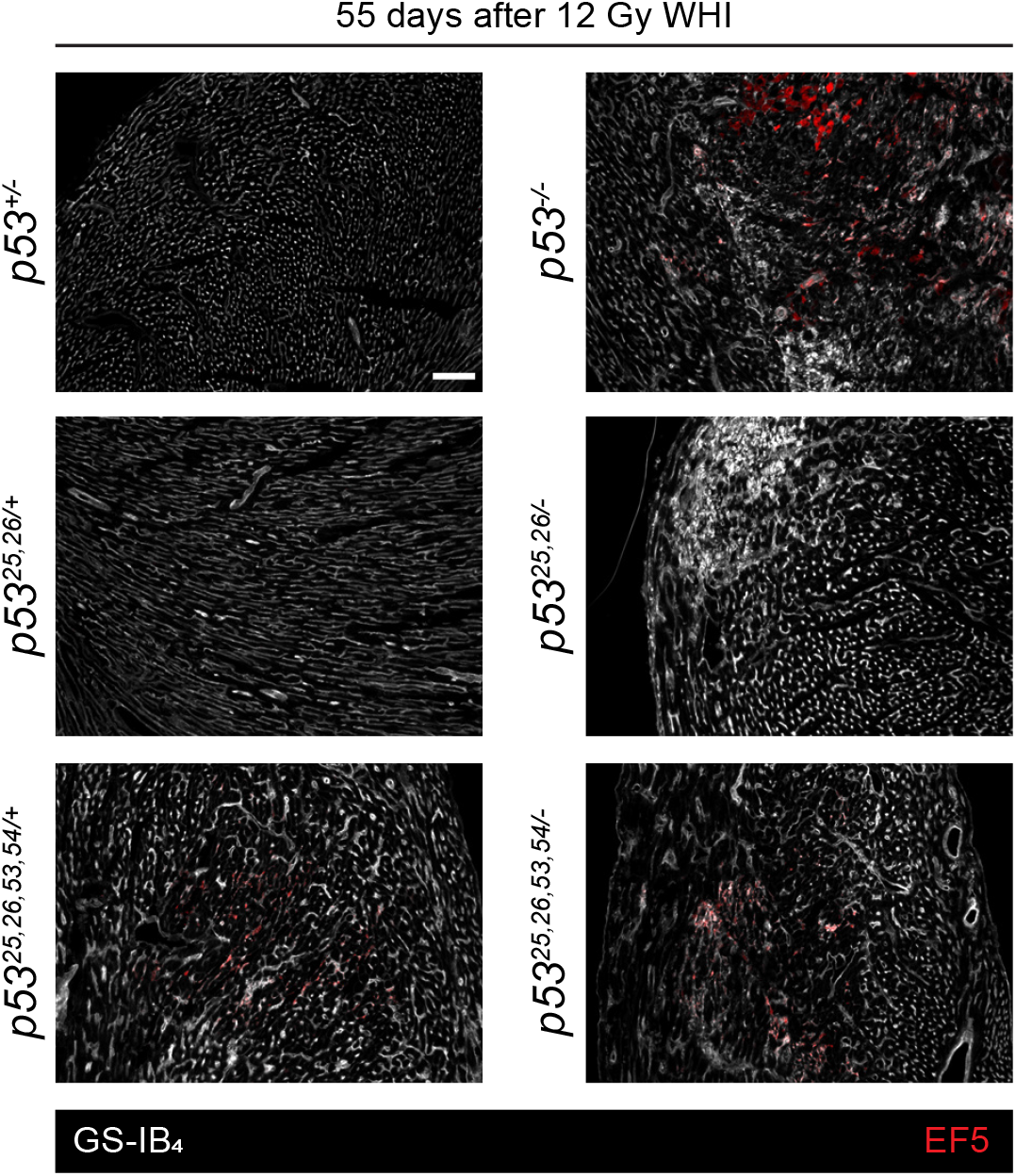
p53 TAD functions in endothelial cells contribute to limit hypoxia and/or a disorganized vasculature in the heart. Representative images of frozen sections of hearts stained with an antibody against EF5 (red) and counterstained with isolectin GS-IB4 (white). The hearts were harvested from mice of different genotypes at day 55 after 12 Gy WHI. EF5 solution (10mM) was administered through intraperitoneal injection at 26.5μL/g body weight to the mice, and the hearts were harvested 3 hours after the injection. Scale bars, 100μm.

## Discussion

In the context of radiation-induced cardiac injury, we showed that both endothelial-specific loss of p53 (*VE-Cre; p53^FL/FL^* mice with p53 null endothelial cells) and complete loss of p53 transactivation ability (p53^25,26,53,54^) sensitized mice to radiation-induced injury to the heart. Compromised p53 TAD1 function (p53^25,26^) also resulted in radiation-induced cardiac injury. This indicates that canonical p53 target genes that rely on functional p53 TAD1 in endothelial cells are critical to protect the heart after WHI. For example, p53^25,26^ does not drive robust radiation-induced *p21* expression after DNA damage (Brady et al., 2011), and p21 is known to protect irradiated hearts from injury (Lee et al., 2012). However, in mice lacking p53 TAD1 function in endothelial cells (*VE-Cre; p53^LSL-25,26/FL^*), the extent of hypoxia and myocardial necrosis after WHI was not as severe as mice with endothelial cells lacking p53 (*VE-Cre; p53^FL/FL^* mice). Therefore, non-canonical targets of p53 in endothelial cells may also contribute to limit the extent of radiation-induced pathological changes in the heart. Dissecting the transcriptomic changes in endothelial cells from irradiated hearts from *VE-Cre; p53^LSL-25,26/FL^* mice and controls that retain wild type p53 in the endothelial cells will be an interesting future direction to pursue. Moreover, comparing the transcriptional programs in endothelial cells from *VE-Cre; p53^LSL-25,26/FL^* mice and *VE-Cre; p53^FL/FL^* mice has the potential to identify candidate genes in addition to *p21* that control DNA damage-induced late effects in general and radiation-induced cardiac injury in particular. While p53^25,26^ fails to induce transcription of many canonical p53 target genes after DNA damage, it still retains the ability to activate a subset of non-canonical p53 target genes and express a basal level of *p21* transcription (Brady et al., 2011). Therefore, it is possible that robust radiation-induced transcription of the non-canonical p53 targets and/or basal expression of *p21* or other genes in endothelial cells function to prevent the most profound pathological changes in the heart after WHI. Regardless of the exact mechanism, our results show that this function of p53^25,26^ fails to prevent radiation-induced cardiac injury. It is possible that the levels of necrosis in the *VE-Cre; p53^LSL-25,26/FL^* mice were sufficient to induce heart failure. However, it is also possible that dysregulated p53 TAD functions might lead to cardiac injury through mechanisms other than necrosis.

One limitation of this study is selection bias. Only the hearts of mice that survived until the EF5 injection on the 55^th^ day following WHI could be subjected to histopathological examination. Some mice of the genotypes that were susceptible to radiation-induced cardiac injury (*VE-Cre; p53^FL/FL^, VE-Cre; p53^LSL-25,26/FL^*, and *VE-Cre; p53^LSL-25,26,53,54/FL^*) died before or right at injection, preventing them from being included in the histological assays. This could lead to underrepresentation of the samples that developed more severe phenotypes. While this would shift the histological findings toward the milder side, this would affect all groups that lost mice prior to day 55. Although it is possible that studying the hearts from mice less than 55 days after WHI might reveal a more severe phenotype, the available data indicate a correlation between an intact p53 transcriptional program in endothelial cells and the control of radiation-induced cardiac injury.

The p53 protein functions by forming tetramers to transactivate target genes. Interestingly, p53^25,26,53,54^ can form tetramers with wild type p53 and the *p53^25,26,53,54/+^* genotype results in upregulation of a subset of p53 target genes compared to *p53^+/+^* wild type only, due to stabilization of wild-type p53 by p53^25,26,53,54^ (Van Nostrand et al., 2014). Moreover, the *p53^25,26/+^* genotype also results in enhanced transactivation of some p53 target genes relative to p53 wild type only (Bowen et al., 2019). This indicates that with both a TAD mutant and wild type p53 in the cell, the transactivation behavior of the p53 proteins should not be directly inferred from the genotype. Rather, abnormal p53 tetramers alter the target gene repertoire. In mouse embryos, many p53 target genes including *p21* and *Noxa* are expressed at significantly higher levels in the *p53^25,26/+^* and *p53^25,26,53,54/+^* genotypes, compared to wild type p53, because of the stabilization of wild-type p53 (Bowen et al., 2019). We found that a single wild type *p53* allele in endothelial cells was sufficient to protect mice from radiation-induced cardiac death, but it possessed a variable capacity to prevent hypoxia/necrosis depending on the presence of specific p53 TAD mutants within the same cell.

In conclusion, we identified a critical role for p53 transactivation functions in controlling radiation-induced late effects and that dysfunctional TAD1 alone in endothelial cells was sufficient to promote the development of cardiac injury following radiation exposure. This highlights the importance of p53 TAD1 function in endothelial cells to protect the heart against radiation-induced cardiac injury.

---

**SUPPLEMENTARY FIG. 1.**
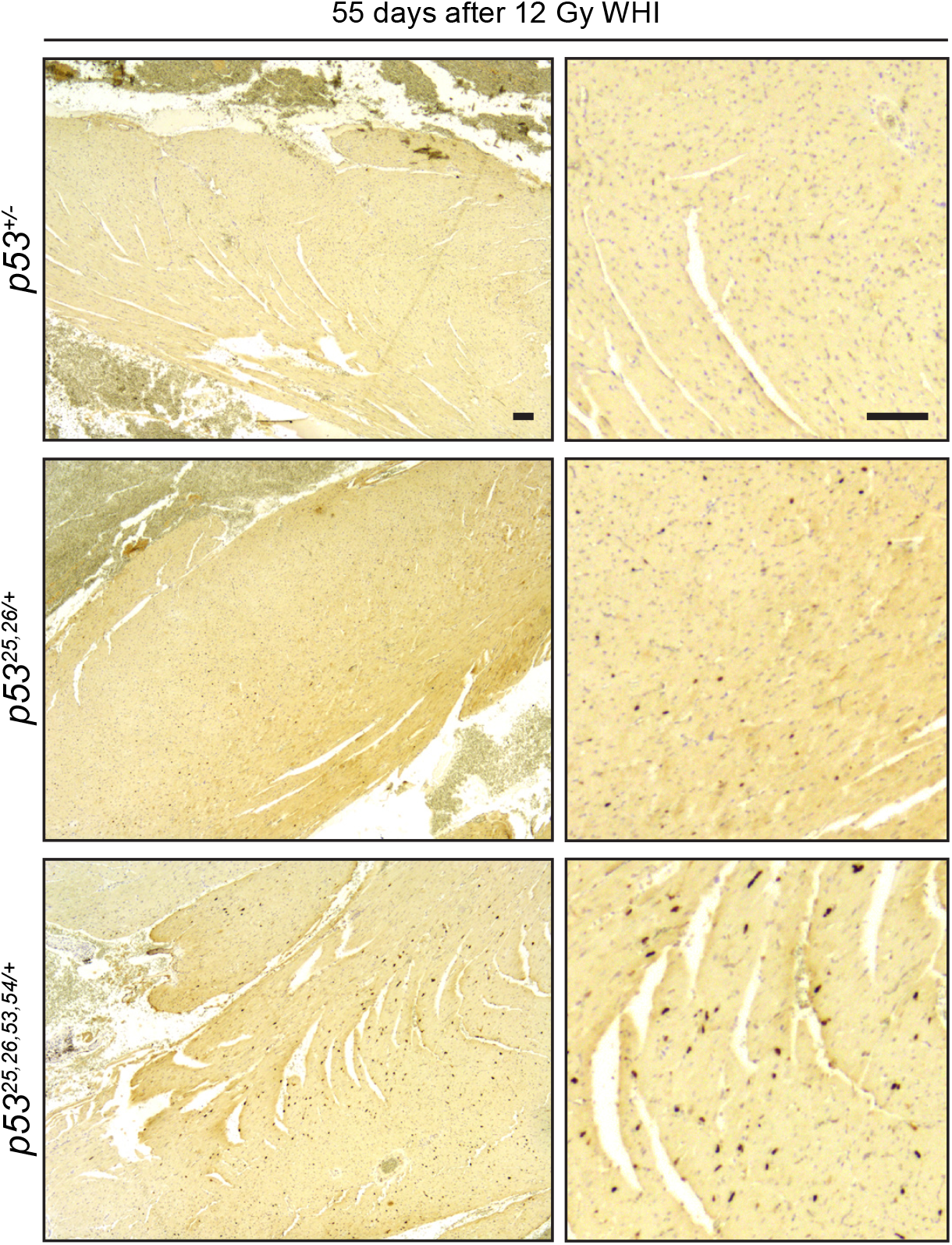
p53 proteins accumulate in endothelial cells of the heart from mice that express p53 TAD mutants. Representative p53 protein immunohistochemistry (IHC) images of hearts harvested from mice of different genotypes at day 55 after 12 Gy WHI. The images are shown at two magnifications. Scale bars, 100μm.

